# Functions of PMS2 and MLH1 important for regulation of divergent repeat-mediated deletions

**DOI:** 10.1101/2024.08.05.606388

**Authors:** Hannah Trost, Felicia Wednesday Lopezcolorado, Arianna Merkell, Jeremy M. Stark

## Abstract

Repeat-mediated deletions (RMDs) are a type of deletion rearrangement that utilizes two repetitive elements to bridge a DNA double-strand break (DSB) that leads to loss of the intervening sequence and one of the repeats. Sequence divergence between repeats causes RMD suppression and indeed this divergence must be resolved in the RMD products. The mismatch repair factor, MLH1, was shown to be critical for both RMD suppression and a polarity of sequence divergence resolution in RMDs. Here, we sought to study the interrelationship between these two aspects of RMD regulation (i.e., RMD suppression and polar divergence resolution), by examining several mutants of MLH1 and its binding partner PMS2. To begin with, we show that PMS2 is also critical for both RMD suppression and polar resolution of sequence divergence in RMD products. Then, with six mutants of the MLH1-PMS2 heterodimer, we found several different patterns: three mutants showed defects in both functions, one mutant showed loss of RMD suppression but not polar divergence resolution, whereas another mutant showed the opposite, and finally one mutant showed loss of RMD suppression but had a complex effect on polar divergence resolution. These findings indicate that RMD suppression vs. polar resolution of sequence divergence are distinct functions of MLH1-PMS2.

**HIGHLIGHTS:**

- MLH1-PMS2 suppresses divergent repeat-mediated deletions (RMDs).
- MLH1-PMS2 promotes polar resolution of sequence divergence.
- Several mutants of MLH1-PMS2 affect both aspects of RMDs.
- Some MLH1-PMS2 mutants affect only one aspect of RMDs.
- Suppression of RMDs vs. polar resolution of divergence appear distinct.

## INTRODUCTION

Repeat-mediated deletions (RMDs) are a type of chromosomal rearrangement that utilizes two repetitive elements to bridge a DNA double-strand break (DSB) [1, 2]. Repetitive elements, such as long interspersed elements and short interspersed elements are frequent in mammalian genomes [3, 4]. One example of such elements is *Alu*-like short interspersed elements which are roughly 300 bp in length, present in more than 1 million copies in the human genome, contain varying amounts of sequence divergence between one another, and are known to cluster in genic regions of the genome [5–7]. RMDs between *Alu* elements have been shown to disrupt tumor suppressor genes and can cause increased risk for various cancers [8, 9]. A proposed mechanism for RMD formation is single-strand annealing (SSA) which requires 5’ end resection on either side of the DNA DSB to reveal the two repeats, allowing the repeats to anneal. This annealing causes the formation of 3’ nonhomologous tails which are removed prior to ligation, resulting in an RMD [10]. Since RMDs are inherently mutagenic, several pathways appear to suppress these events [1, 2, 10–12]. In particular, RMDs between divergent repeats are prone to heteroduplex rejection mechanisms via the DNA mismatch repair (MMR) pathway, and the BLM/TOP3a/ RMI1/RMI2 complex [13–16].

As a brief summary of MMR, mismatches are recognized by MSH2-MSH6 or MSH2- MSH3, which upon recognition recruits the predominant MLH1 heterodimer MLH1-PMS2 (PMS2 in mammals is PMS1 in *S. cerevisiae*; for simplicity we will use PMS2) [17–20]. The MLH1-PMS2 heterodimer has endonuclease activity enabling it to create a nick on the newly synthesized DNA strand upstream of the mismatch [21–24]. This nick allows for excision through various pathways and proper fill-in resulting in increased fidelity of replication [25–28]. In bacteria, the newly synthesized DNA strand lacks methylation which serves as the strand discrimination signal for mismatch repair to create nicks to excise the newly synthesized strand as compared to the template strand [29]. The mechanism for this strand discrimination in mammalian cells remains unclear [30, 31]. It has been proposed from biochemical assays that a nick at the 3’ edge of the replication fork, as would occur with okazaki fragments, is the strand discrimination signal allowing for proper template stranded base retention [17, 31, 32]. As mismatch repair is associated with the presence of the replication fork, the orientation of PCNA and the presence of this nick at the edge of the replication fork has the potential to provide such strand discrimination signals to MSH2-MSH6 and MLH1-PMS2 [21, 31, 33].

In RMDs, the presence of mismatched bases between repeats causes suppression by both MSH2-MSH6 and MLH1-PMS2 [2, 11, 12]. Indeed, for RMD suppression between divergent repeats MSH2 and MLH1 act in the same pathway [11]. A recent study showed that MMR proteins are not only involved in suppression of the divergent RMDs, but also affect the resolution of the mismatches in the events that escape heteroduplex rejection [11]. Specifically, the mismatches in RMDs are resolved in a way that leads to a strand polarity in the products where the mismatches closest to the DNA DSB ends are removed and filled in to match the annealed strand, which is consistent with 3’ end-directed strand discrimination [11]. Notably, this polarity was dependent on the C-terminal residues of MLH1 that contribute to the MLH1-PMS2 endonuclease domain [11]. Therefore, MLH1 suppresses divergent RMDs while also mediating mismatch resolution in the RMD products for the events that escape MMR suppression. In this study, we sought to examine the relationship between these two aspects of RMD regulation via MLH1-PMS2 (suppression of the events vs. polar resolution of the divergent bases) by examining effects of PMS2 loss, along with six mutants of the MLH1-PMS2 heterodimer.

## RESULTS

### PMS2-E702K and PMS2-721-4A (PMS2-721QRLITP > 721ARAAAP) fail to suppress **divergent RMDs.**

To examine roles of PMS2 in regulation of divergent RMDs, we examined *Pms2^-/-^* cells, and also tested two PMS2 mutants using complementation vectors. For this, we used a previously described reporter system (3%RMD-GFP, Fig 1A) [1]. This reporter is integrated into chromosome 17 of mouse embryonic stem cells (mESCs) and utilizes two tandem 287 bp repeats (represented as “R”) separated by 0.4 Mbp. The 5’ repeat is endogenous sequence downstream of the *Cdkn1A* promoter, and the 3’ repeat is fused to GFP and targeted to the *Pim1* locus. The 3’ repeat contains 8 equally spaced mismatches relative to the 5’ repeat for 3% sequence divergence. An RMD event utilizing the two repeats leads to a *Cdkn1A-GFP* fusion gene that causes GFP+ cells which can then be measured by flow cytometry. To induce an RMD using this reporter two Cas9/sgRNAs are used; the 5’ DSB at 268 bp downstream of the 5’ repeat, and the 3’ DSB at 16 bp upstream of the 3’ repeat. All assay conditions using this reporter are normalized to transfection efficiency with parallel transfections with a GFP expression vector.

**Fig 1.**
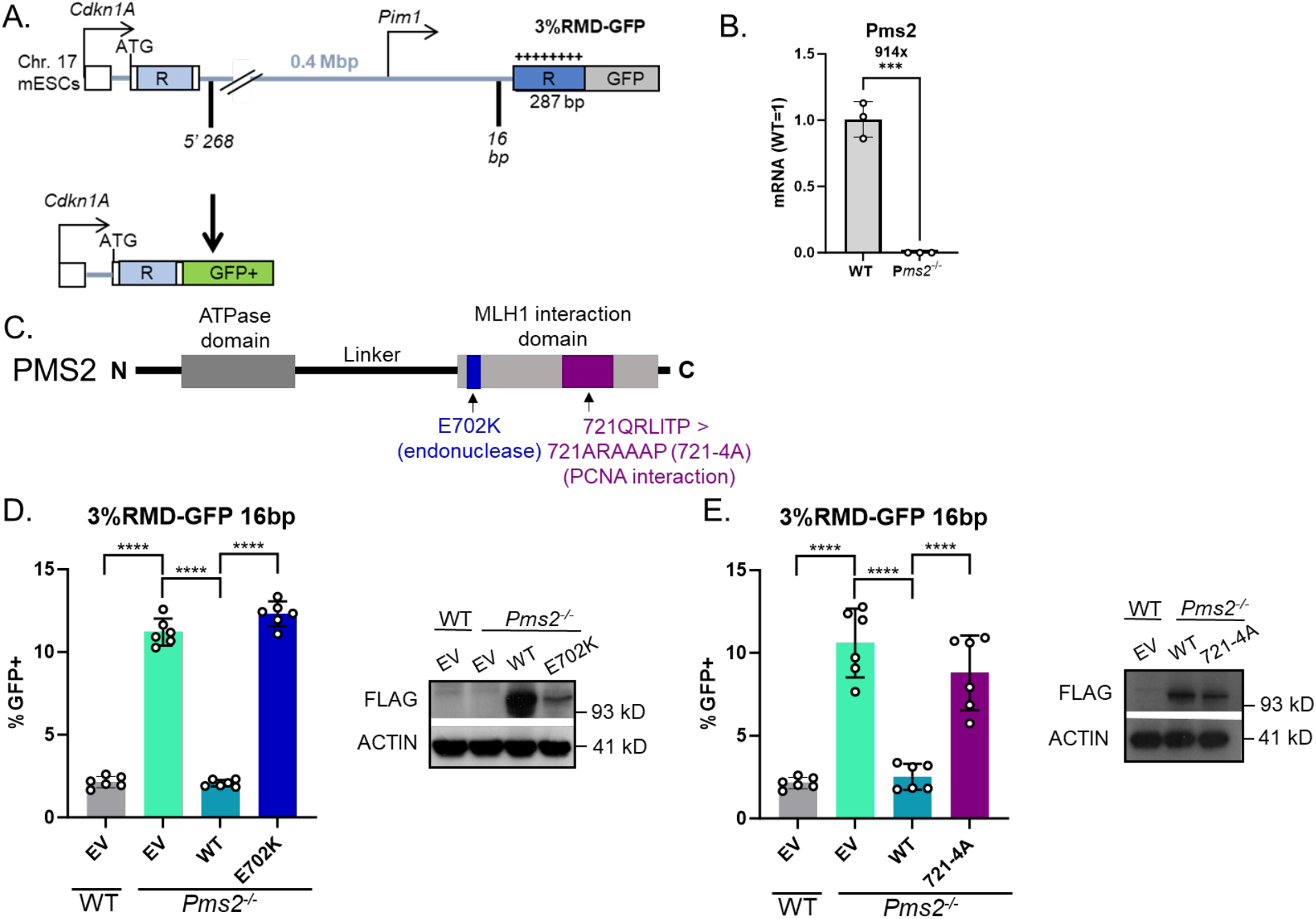
PMS2-E702K and PMS2-721-4A (PMS2-721QRLITP > 721ARAAAP) fail to suppress divergent RMDs. **(A)** Shown is the 3%RMD-GFP reporter, which is integrated into the *Pim1* locus in chromosome 17 of mESCs. The two repeats are denoted as “R”. The 5’ repeat is endogenous sequence. The 3’ repeat is fused to GFP and contains 3% sequence divergence (8 equally spaced mismatches) relative to the 5’ repeat. RMDs are induced by two DSBs: one at 268 bp downstream of the 5’ repeat, the other at 16 bp upstream of the 3’ repeat, such that repair of the two DSBs by an RMD leads to GFP+ cells. **(B)** Shown is qRT-PCR analysis of PMS2 in WT mESCs and the *Pms2^-/-^* mESCs. Shown is the mRNA abundance of PMS2 based on threshold cycle (Ct) values from PCR amplification, normalized to ACTIN, relative to WT cells (WT=1). n=3 PCR. ***p ≤ 0.0005, unpaired *t*-test. **(C)** Schematic diagram of PMS2 noting the location of E702K and 721-4A mutations, based on the references described in the text. **(D)** Shown are the RMD frequencies for the reporter shown in (A) in *Pms2^-/-^* mESCs transfected with EV, PMS2- WT, or PMS2-E702K. Frequencies are normalized to transfection efficiency. n=6. ****p < 0.0001, multiple unpaired *t*-tests with Holm-Sidak correction. Immunoblot show levels of FLAG-tagged PMS2 and E202K mutant. **(E)** Shown are the RMD frequencies in *Pms2^-/-^* mESCs transfected with EV, PMS2-WT, or PMS2-721-4A. Frequencies are normalized to transfection efficiency. n=6. ****p < 0.0001, unpaired *t*-tests. Immunoblot show levels of FLAG-tagged PMS2 and 721-4A mutant. Data are represented as mean values ± SD.

A prior study showed that both components of the MLH1-PMS2 heterodimer have a role in suppressing divergent RMDs [11]. Therefore, we sought to identify how two mutants of PMS2 affected its role in suppressing these events. The prior study used PMS2 RNAi approaches, and so for this study, we first generated a *Pms2^-/-^* mESC line by targeting sgRNAs/Cas9 to exon 11 of *Pms2* that we confirmed had loss of PMS2 transcript by qRT-PCR (Fig 1B). We compared the results of the RMD assays in the *Pms2^-/-^* cell line vs. WT, and *Pms2^-/-^* cells transfected with various PMS2 expression vectors (Fig 1C). Beginning with the comparison of *Pms2^-/-^* vs. WT mESCs, we found that loss of PMS2 caused a significant increase in the frequency of divergent RMDs, consistent with prior findings with RNAi depletion (Fig 1D) [11].

We then examined effects of expressing PMS2 WT and two mutants (Fig 1C) [34–36]. For the first mutant of PMS2, we tested a portion of the C-terminus of PMS2 that contributes to the MLH1-PMS2 nuclease domain. Specifically, we tested a PMS2 endonuclease deficient mutant (E702K) expression vector, which disrupts the metal binding domain of the MLH1-PMS2 heterodimer [34]. We examined the frequency of RMD events in *Pms2^-/-^* mESCs transfected with EV, WT-PMS2 or E702K and found that expression of WT-PMS2, but not E702K, cause a significant decrease in RMD frequency (Fig 1D). We next examined the PMS2-721QRLITP > 721ARAAAP (721-4A) mutant [35, 36]. Biochemical data using purified proteins suggests that PCNA stimulates MLH1-PMS2 endonuclease activity, and that the PMS2-721-4A mutant disrupts the interaction between PMS2 and PCNA [35, 36]. We evaluated the RMD frequency in *Pms2^-/-^* mESCs transfected with EV, WT-PMS2 or 721-4A and found that expression of WT-PMS2, but not 721-4A, cause a significant decrease in RMD frequency (Fig 1E). We confirmed expression of the PMS2 vectors via immunoblotting using a 3xFLAG immunotag (Fig 1D, 1E). Notably each mutant was expressed somewhat lower than WT. The lower expression of these defective PMS2 mutants is consistent with reports that disease-associated alleles of MMR factors (e.g., MLH1) are often expressed below WT levels [37]. In summary, these findings indicate that the endonuclease domain and interaction with PCNA by PMS2 are required for the ability of PMS2 to suppress RMDs between divergent repeats. Although, of course these mutants could affect other functions of PMS2.

### PMS2-E702K fails to promote polar resolution of sequence divergence during RMDs, whereas PMS2-721-4A is proficient

We next considered whether the two PMS2 mutants influenced the polar resolution of sequence divergence in the RMD product. To test this, we used the same reporter described above (3%RMD-GFP) and examined whether the top or bottom base in each of the 8 mismatches was retained in the RMD products (Fig 2A). We performed the reporter assay as described in WT mESCs, *Pms2^-/-^* mESCs, and *Pms2^-/-^* mESCs complemented with WT-PMS2, E702K, or 721-4A expression vectors. We sorted for the GFP+ cells for each condition by flow cytometry, amplified the across the retained repeat in the fusion gene, and performed deep sequencing analysis. We then scored each of the 8 mismatches in each read for having retained the base from the 5’ repeat from the *Cdkn1A* locus (top strand, blue triangles), or from the 3’ repeat that was fused to GFP (bottom strand, pink triangles) (Fig 2A). We performed this analysis using three independent transfections/sorts for each of the conditions to determine the mean/standard deviation for each condition for the frequency of retention of the top stranded base at each mismatch.

**Fig 2.**
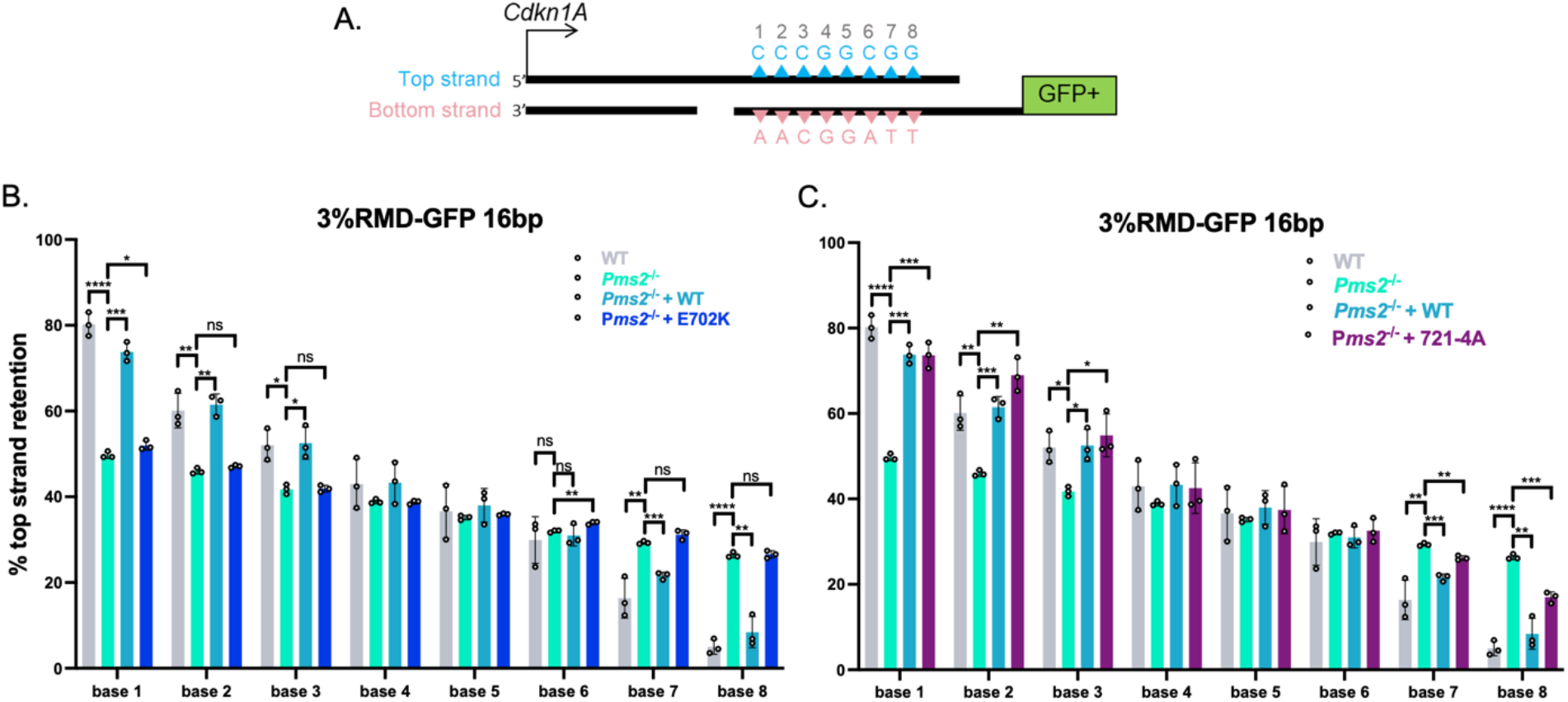
PMS2-E702K fails to promote polar resolution of sequence divergence during RMDs, whereas PMS2-721-4A is proficient. **(A)** Shown is a diagram of the RMD annealing intermediate in the 3%RMD-GFP reporter. Triangles represent the 8 mismatches of the top and bottom strand, numbered 1-8 with mismatched bases shown above and below the triangles. The DSB ends are shown without 3’ non-homologous tails for simplicity. **(B)** Mismatch resolution was examined by sorting the GFP+ cells, PCR amplifying the repeat sequence, and performing deep sequencing analysis to determine the amount of top strand retention for WT, and *Pms2^-/-^* mESCs transfected with expression vectors for WT and E702K. n=3. *p ≤ 0.05, **p ≤ 0.005, ***p ≤ 0.0005, ****p < 0.0001, ns = not significant. WT vs. *Pms2^-/-^*, unpaired *t*-test, Pms*2^-/-^* vs. *Pms2^-/-^* + PMS2 (WT) and *Pms2^-/-^* + E702K, unpaired *t*-test with Holm-Sidak correction. **(C)** Frequency of top strand base retention at 16 bp in WT and *Pms2^-/-^* mESCs transfected with expression vectors for WT and + 721-4A. WT, *Pms2^-/-^* , and *Pms2^-/-^*+ PMS2 (WT) are the same as in (B). n=3. *p ≤ 0.05, **p ≤ 0.005, ***p ≤ 0.0005, ****p < 0.0001, ns = not significant. WT vs. *Pms2^-/-^*, unpaired *t*-test, Pms*2^-/-^* vs. *Pms2^-/-^*+ PMS2 (WT) and *Pms2^-/-^* + 721-4A, unpaired *t*-test with Holm-Sidak correction. Data are represented as mean values ± SD.

We found that retention of the top stranded base showed a polarity in WT mESCs, confirming previously published results [11]. Specifically, at base 1 (the base closest to *Cdkn1A*) there is a preferential retention of the top strand base (Fig 2B). Conversely, at base 8 (the base closest to GFP) there is a preferential retention of the bottom strand base (Fig 2B). The bases in the middle show no strong bias towards top or bottom base (Fig 2B). This strand polarity in the mismatch resolution is consistent with preferential loss of the mismatches closes to the DSB end in the SSA model for RMDs, and hence 3’end-directed strand discrimination (Fig 2A).

We then examined this pattern in *Pms2^-/-^* mESCs, and *Pms2^-/-^* mESCs expressing PMS2- WT, E702K, or 721-4A expression vectors. First, we found that in *Pms2^-/-^* mESCs the strand polarity is largely lost (Fig 2B). Specifically, at bases 1-3 the *Pms2^-/-^*cells showed a significant decrease in bias towards maintaining the top strand base compared to WT cells, and at bases 7 and 8 showed a significant increase in bias towards keeping the top stranded base compared to WT cells. The middle bases maintained the pattern of WT cells, showing no preference towards top or bottom stranded base. While the mismatch resolution polarity is still detectible (base 1 still maintains more top stranded base than base 8) in the *Pms2^-/-^* cells, the polarity is markedly reduced compared to WT cells.

Next, with the PMS2 expression vectors, we found that expression of PMS2-WT, but not E702K, restored the strand polarity back to WT, specifically increasing top strand base retention at bases 1-3, and conversely reducing top strand retention at bases 7 and 8. We also performed this assay for strand polarity in *Pms2^-/-^*mESCs complemented with the 721-4A mutant (Fig 2C). We found that at bases 1-3, expression of the 721-4A mutant restored the strand polarity, specifically increasing top strand base retention. Similarly, at bases 7 and 8 expression of 721-4A reduced top strand retention vs. the empty vector (EV) control, albeit not fully restoring the frequencies to WT mESC levels. In summary, we found PMS2-E702K causes loss of the polarity of mismatch resolution in RMD products whereas PMS2-721-4A supports this aspect of RMD regulation. Based on the known biochemical effects of these mutants [34, 35], these findings indicate that the nuclease domain of MLH1-PMS2 is important for this polarity in the RMD products, whereas the interaction of PMS2 with PCNA, via the 721QRLITP motif, is dispensable. Although, as mentioned above, of course these mutants could also be affecting other aspects of MLH1-PMS2 function.

### MLH1-KQQ (K57C, Q60L, Q62L) and MLH1-566-3A (566QILIYDF>AILAYDA) fail to **suppress divergent RMDs.**

MLH1 has several proposed functions that we posited might be important for MLH1-PMS2 to suppress divergent RMDs, which informed our choice of testing four mutants. The first mutant was based on studies of MLH1 interactions with MSH2-MSH6 [11]. Specifically, we tested if mutating residues in MLH1 implicated in the interaction between the two heterodimers would affect the ability of MLH1 to suppress RMDs (MLH1 K57C, Q60L, Q62L, MLH1-KQQ) [38]. For the second mutant, we tested residues in MLH1 proposed to promote an interaction of the MLH1-PMS2 heterodimer with PCNA (MLH1 566QILIYDF>AILAYDA, MLH1-566-3A) [39], although another study found that this mutant of MLH1 retained robust PCNA-activated nuclease activity [35]. To test these mutants, we generated mutant MLH1 expression vectors (Fig 3A). We found that WT-MLH1 was able to suppress divergent RMDs, however both the KQQ mutant and the 566-3A mutant failed to suppress divergent RMDs (Fig 3B). We confirmed expression of both mutant proteins via immunoblot, although not to the same level as the WT protein (Fig 3C), which as mentioned above is consistent with studies that deficient MLH1 alleles are often expressed below WT levels [37].

**Fig 3.**
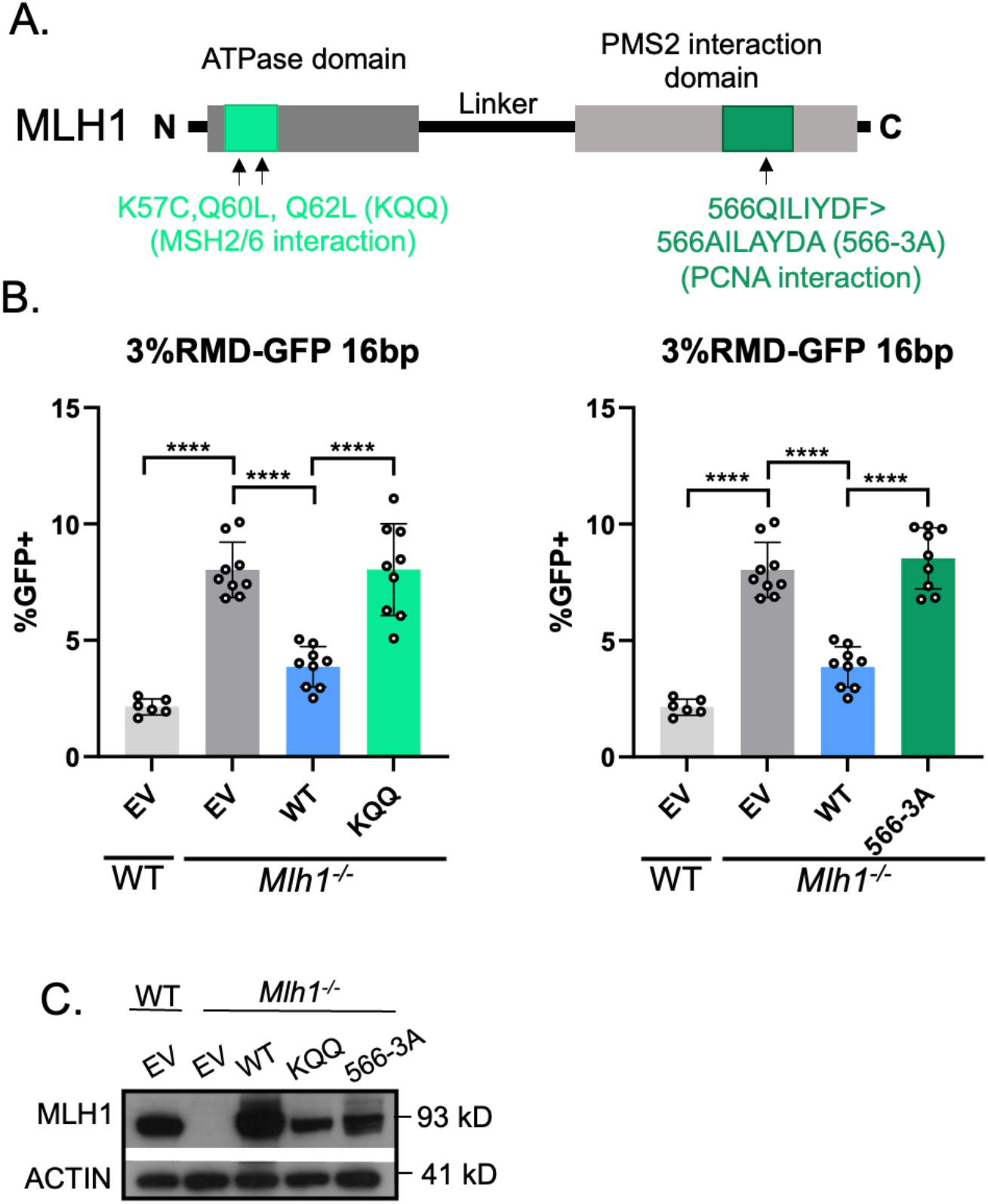
MLH1-KQQ and MLH1-566-3A fail to suppress divergent RMDs. **(A)** Schematic diagram of MLH1 noting the location of KQQ and 566-3A mutations. **(B)** Shown are the RMD frequencies in *Mlh1^-/-^* mESCs transfected with EV, MLH1-WT, MLH1-KQQ, or MLH1-566-3A. Frequencies are normalized to transfection efficiency. n=6 for WT, n=9 for *Mlh1^-/-^*+ WT, KQQ, and 566-3A. ****p < 0.0001, unpaired *t*-tests. **(C)** Immunoblot show levels of MLH1-WT, KQQ, and 566-3A. Data are represented as mean values ± SD.

### MLH1-KQQ fails to promote polar resolution of sequence divergence at bases 1 and 2, but not 7 and 8, whereas MLH1-566-3A shows a defect at each base

We next tested whether the MLH1-KQQ and MLH1-566-3A mutants affect the resolution of the RMD product, using the method as described above (Fig 2). We first confirmed prior findings that loss of MLH1 leads to a striking loss of strand polarity which is restored by expression of MLH1-WT [11]. Specifically, we found that bases 1 and 2 showed a strong bias in maintaining the top stranded base, and bases 7 and 8 showed a strong bias towards maintaining the bottom stranded base in WT cells, which is lost in the *Mlh1^-/-^* cells, and subsequently restored in the *Mlh1^-/-^* cells transfected with MLH1-WT (Fig 4A).

**Fig 4.**
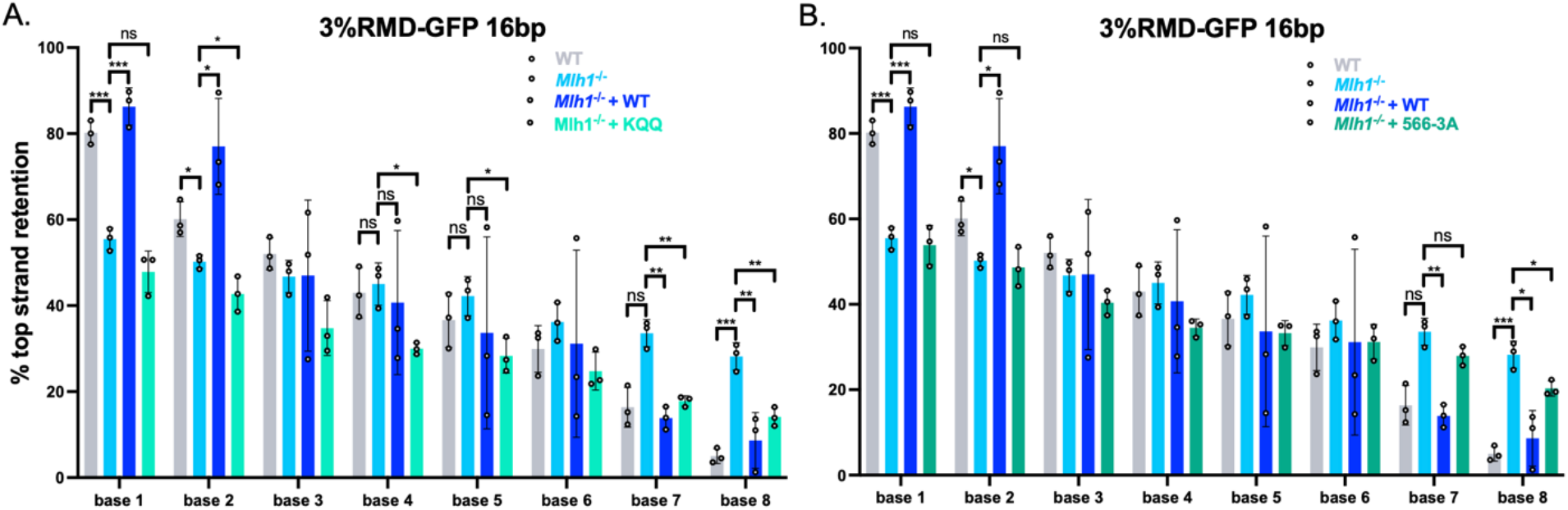
MLH1-KQQ fails to promote polar resolution of sequence divergence at bases 1 and 2, but not 7 and 8, whereas MLH1-566-3A shows a defect at each base. **(A)** Shown is the frequency of top strand base retention for WT, *Mlh1^-/-^*, *Mlh1^-/-^* + MLH1 (WT), *Mlh1^-/-^* + KQQ mESCs. WT samples as in Fig 2B. n=3. *p ≤ 0.05, **p ≤ 0.005, *** ≤ 0.001. WT vs. *Mlh1^-/-^*, unpaired *t*-test, *Mlh1^-/-^* vs. *Mlh1^-/-^* + MLH1 (WT) and *Mlh1^-/-^* + KQQ, unpaired *t-*tests with Holm- Sidak correction. **(B)** Frequency of top strand base retention for WT, *Mlh1^-/-^*, *Mlh1^-/-^* + MLH1 (WT), *Mlh1^-/-^* + 566-3A mESCs. WT samples as in Fig 2B. n=3. *p ≤ 0.05, **p ≤ 0.005, *** ≤ 0.001. WT vs. *Mlh1^-/-^*, unpaired *t*-test, *Mlh1^-/-^* vs. *Mlh1^-/-^* + MLH1 (WT) and *Mlh1^-/-^* + 566-3A, unpaired *t-*tests with Holm-Sidak correction. Data are represented as mean values ± SD.

In contrast, the MLH1-KQQ mutant showed a complex pattern of sequence divergence resolution (Fig 4A). Specifically, at bases 1 and 2 the MLH1-KQQ mutant, failed to promote top strand base retention (i.e., was similar to *Mlh1^-/-^*cells). In contrast, at bases 7 and 8 the KQQ mutant was proficient at suppressing top strand base retention (i.e., was significantly lower than *Mlh1^-/-^*, and hence similar to the MLH1-WT complemented condition). In comparison, the MLH1- 566-3A mutant failed both to promote top strand base retention at bases 1 and 2, as well as suppress top strand base retention at base 7 and 8, although at base 8, top strand bias was statistically lower than the empty vector control (Fig 4B). In summary, the MLH1-KQQ mutant fails to suppress RMDs, and retains activity for promoting polarity of divergence resolution, but only on one side of the events. In contrast, the MLH1-566-3A mutant showed defects in both aspects of RMD regulation.

### MLH1-RADA (R385A, D387A) fails to suppress divergent RMDs, whereas MLH1-E34A is proficient

We then examined two more mutants of MLH1 [40, 41] (Fig 5A). The first mutant, MLH1- E34A, is proposed to disrupt the function of the ATPase domain [40]. Interestingly, the ATPase domain is a critical region of human MLH1, in that a majority of mutations leading to HNPCC kindreds are found in or around this region [42–44]. For the second mutant, we tested residues in the unstructured linker domain of MLH1 (MLH1-R385A, D387A, MLH1-RADA), which is proposed to be required for endonuclease function and is also mutated in cancers [41, 45]. We found that E34A suppressed divergent RMDs to the same level of WT-MLH1, however RADA failed to suppress divergent RMDs (Fig 5B). We confirmed expression of both mutant proteins via immunoblot, which for these mutants were similar to WT levels (Fig 5C).

**Fig 5.**
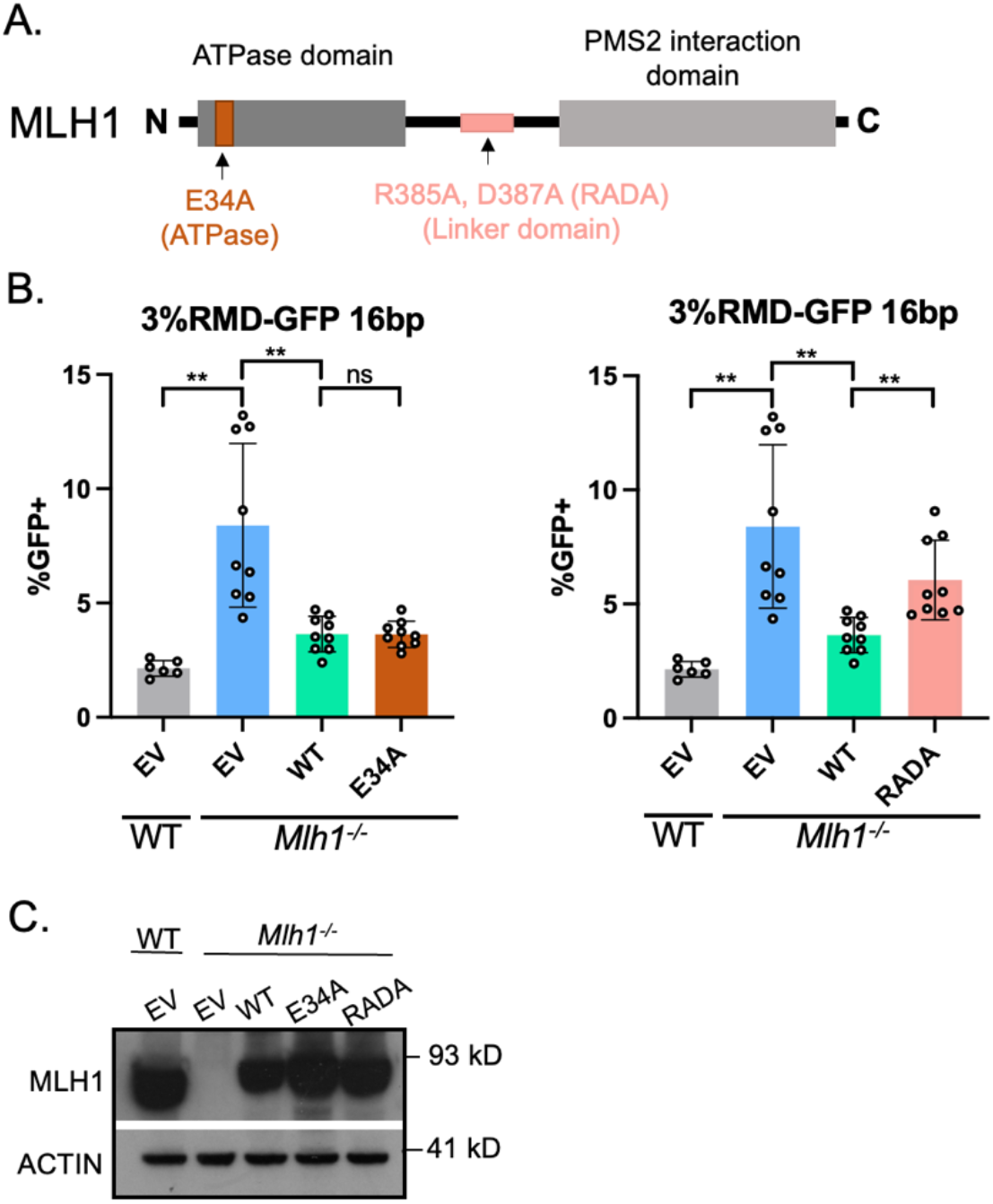
MLH1-E34A, but not MLH1-RADA, suppresses divergent RMDs. **(A)** Schematic diagram of MLH1 noting the location of E34A and RADA mutations. **(B)** Shown are the RMD frequencies in *Mlh1^-/-^*mESCs transfected with EV, MLH1-WT, MLH1-E34A, or MLH1-RADA. Frequencies are normalized to transfection efficiency. n=6 for WT, n=9 for *Mlh1^-/-^* + WT, E34A, and RADA. **p < 0.005, ns = not significant, unpaired *t*-tests. **(C)** Immunoblot show levels of MLH1-WT, E34A, and RADA. Data are represented as mean values ± SD.

### MLH1-E34A and MLH1-RADA fail to promote polar resolution of sequence divergence during RMDs

We next tested whether the MLH1-E34A and MLH1-RADA mutants affect the resolution of the RMD products, using the assays as described above (Figs 2, 4). Interestingly, we found that MLH1-E34A and MLH1-RADA were defective in promoting polar resolution of sequence divergence (Fig 6A, 6B). Specifically, both E34A and RADA failed to promote top strand base retention at bases 1 and 2, and conversely failed to promote botton strand retention at bases 7 and 8 (i.e., showed similar frequencies to to *Mlh1^-/-^* cells). In summary, MLH1-RADA failed to both suppress divergent RMDs and promote polar resolution of sequence divergence. In contrast, MLH1-E34A promoted suppression of divergent RMDs, but failed to promote polar resolution of sequence divergence.

**Fig 6.**
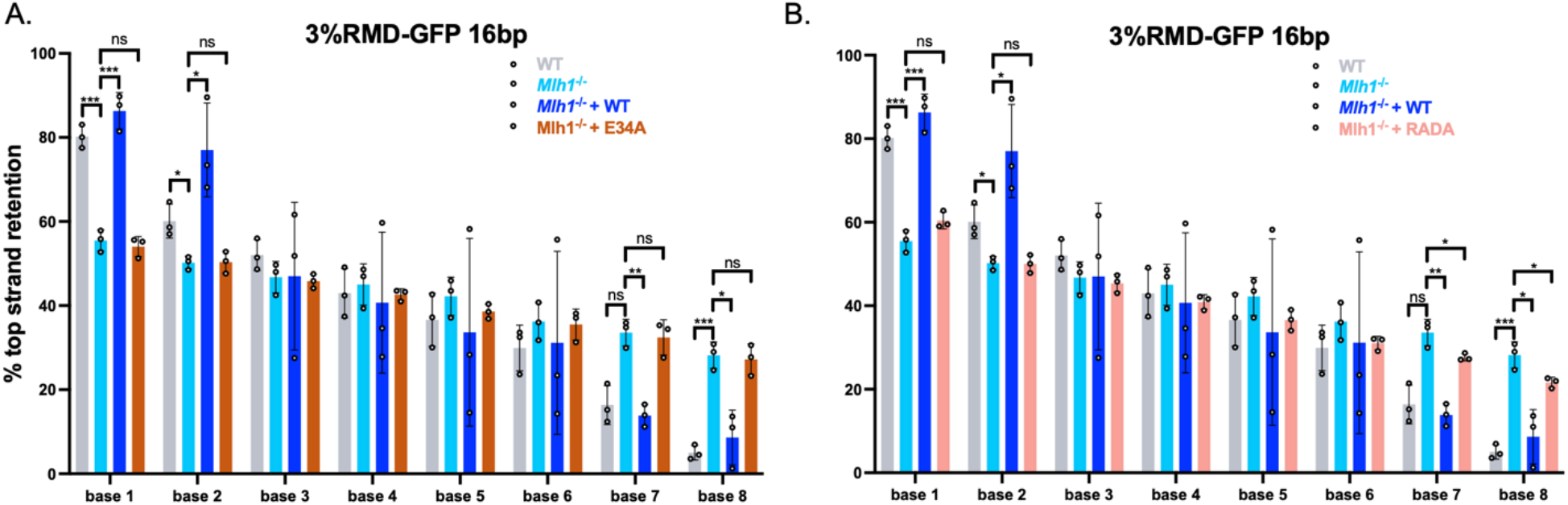
MLH1-E34A and MLH1-RADA fail to promote polar resolution of sequence divergence during RMDs. **(A)** Shown is the frequency of top strand base retention for WT, *Mlh1^/-^*, *Mlh1^-/-^* + MLH1 (WT), *Mlh1^-/-^* + E34A mESCs. WT samples as in Fig 2B. n=3. *p ≤ 0.05, **p ≤ 0.005, p*** ≤ 0.001. WT vs. *Mlh1^-/-^*, unpaired *t*-test, *Mlh1^-/-^* vs. *Mlh1^-/-^* + MLH1 (WT) and *Mlh1^-/-^* + E34A, unpaired *t-*tests with Holm-Sidak correction. **(B)** Frequency of top strand base retention for WT, *Mlh1^-/-^*, *Mlh1^-/-^* + MLH1 (WT), *Mlh1^-/-^* + RADA mESCs. WT samples as in Fig 2B. n=3. *p ≤ 0.05, **p ≤ 0.005, *** ≤ 0.001. WT vs. *Mlh1^-/-^*, unpaired *t*-test, *Mlh1^-/-^* vs. *Mlh1^-/-^*+ MLH1 (WT) and *Mlh1^-/-^* + RADA, unpaired *t-*tests with Holm-Sidak correction. Data are represented as mean values ± SD.

## DISCUSSION

We sought to identify how distinct mutants of PMS2 and MLH1 affect RMD suppression and the polarity of mismatch resolution between divergent repeats. We performed this analysis both to understand the interrelationship between these two aspects of RMD regulation, and also define specific activities of MLH1-PMS2 important for these functions. First, we determined that a *Pms2^-/-^* line showed similar phenotypes as a prior study with *Mlh1^-/-^* (i.e., showed loss of both aspects of RMD regulation) [11], and then we examined six mutants of MLH1-PMS2. Three mutants showed defects in both aspects of RMD regulation: PMS2-E702K, MLH1-566-3A, and MLH1-RADA. In contrast, PMS2-721-4A fails to suppress RMDs between divergent repeats but is sufficient to promote polar resolution of sequence divergence. Conversely, MLH1-E34A mutant interestingly was able to suppress divergent RMDs, but failed to promote polar resolution of sequence divergence. Finally, the MLH1-KQQ mutant showed intermediate phenotypes where it failed to suppress divergent RMDs and was partially defective in promoting polar mismatch resolution in the RMD products. Specifically, this mutant failed to support polar resolution of sequence divergence on the 5’ side, but retained this function on the 3’ side. Overall, these findings indicate that suppression of RMD frequencies and polar resolution of the RMD product share some, but not all requirements for MLH1-PMS2 activities, such that these two aspects of RMD regulation appear distinct.

While each individual mutant may affect several aspects of MLH1-PMS2 function, including overall protein stability, we chose these mutants based on published studies on specific activities, such as nuclease activity and interaction with PCNA and MSH2/6, and within conserved motifs. Several of these studies used the *S. cerevisiae* or human proteins, but we refer to the equivalent mutants in mouse, which were used here. To begin with, the PMS2-E702K mutant lacks endonuclease function [34], and we have found that this mutant failed to mediate both aspects of RMD regulation. Similarly, a prior report showed that an MLH1 mutant that disrupts endonuclease active site of MLH1/PMS2 (MLH1-Δ754-756) also failed to support these aspects of RMD regulation [11]. Finally, we also found similar results with another mutant: MLH1-RADA, which is in the conserved linker motif. While the precise role of the linker is unclear, studies in *S. cerevisiae* show this linker is critical for MMR and MLH1 endonuclease activity [41, 46–48]. Thus, altogether, these findings indicate that the MLH1-PMS2 endonuclease activity is important for both aspects of RMD regulation.

We speculate that the MLH1-PMS2 nuclease suppresses RMDs by causing iterative nicking at the multiple mismatches in the divergent RMD annealing intermediate, which destroys the intermediate, thereby blocking the RMD. In contrast, for a subset of events, MLH1-PMS2 only nicks upstream of mismatches that are proximal to the 3’ end / DSB end, which allows for excision and fill-in synthesis, leading to the RMD with polar resolution of the mismatches. These proposed mechanisms are consistent with biochemical studies showing 3’ nick-directed activation of MLH1- PMS2 endonuclease activity on the same strand as the 3’ nick, which is also the proposed polarity for strand discrimination [21–24].

Such strand discrimination is promoted by PCNA, RFC, and MSH2/MSH6, although MLH1-PMS2 endonuclease activity *per se* is not absolutely dependent on these factors [21, 23, 36, 49–51]. Based on these studies, we examined mutants of MLH1-PMS2 that have been shown to disrupt interactions between PCNA and MSH2/MSH6, but our findings do not support a simple model. For example, with the mutants that are implicated in the interaction between MLH1-PMS2 and PCNA, we found distinct results with the PMS2 vs. MLH1 mutant. The MLH1 mutant (i.e. 566-3A) showed near complete loss of function in RMD regulation. However, the influence of this mutant on PCNA-mediated activation of MLH1-PMS2 is controversial [35, 39], and so it may be difficult to relate these phenotypes directly to interaction with PCNA. In contrast, the mutant in PMS2 (721-4A) showed complete loss of RMD suppression while retaining activity to promote polar resolution of sequence divergence. Accordingly, it is possible that the interaction between MLH1-PMS2 with PCNA is not required for polar resolution of divergent sequences, but of course there could be other interactions interfaces with PCNA that remain undefined. Structural information of the complex of MLH1-PMS2 with PCNA/RFC and MSH2/MSH6 would provide insight into this mechanism.

Regarding the role of the interaction between MLH1 and MSH2/MSH6, we examined the MLH1-KQQ mutant that has been shown to disrupt the interaction with MSH2/MSH6 [38]. We found the MLH1-KQQ mutant failed to suppress divergent RMDs, but has a complex phenotype for polar resolution of divergence. In a prior study, MLH1 and MSH2 were both shown to be required for RMD suppression [11], and therefore the MLH1-KQQ mutant failing to suppress divergent RMDs is consistent with these findings. Regarding strand discrimination, MSH2 loss was shown in this prior study to have only modest effects on polarity of mismatch resolution [11]. In contrast, as mentioned above, the MLH1-KQQ mutant shows a complex phenotype: retention of polar resolution of sequence divergence at bases 7 and 8, but loss of this function at bases 1 and 2. The mechanism is unclear for the discrepancy. However, we note that the length of the non-homologous tails are different between the two sides of the likely annealing intermediate. Namely, the bottom strand / 3’ end is 16 bp from the edge of the repeat, whereas the top strand 5’ DSB has 287 bp of non-homologous sequence. It is unclear how the length of the non-homologous tail may affect polar resolution of sequence divergence, but notably in yeast a 3’ non-homologous tail is important for MMR engagement in divergent recombination events [52, 53]. In any case, these findings indicate that the MLH1-KQQ motif is not absolutely required for polar resolution of sequence divergence.

The final MLH1 mutant is MLH1-E34A, which has been shown to disrupt the ATP hydrolysis activity of MLH1 [40, 54]. Furthermore, *in vitro* studies using purified human MLH1 and PMS2 proteins found that MMR activity was significantly decreased with MLH1-E34A [40], and mutations in the ATPase domain of MLH1 are associated with cancer risk [42–44]. Here, we show that MLH1-E34A suppresses divergent RMDs at similar levels as MLH1-WT. However, MLH1-E34A fails to promote polar resolution of sequence divergence. Thus, ATP hydrolysis by MLH1 appears required for polar resolution of sequence divergence, but not RMD suppression. Since polar resolution of sequence divergence is consistent with the polarity of mismatch strand discrimination, we speculate that ATP hydrolysis by MLH1 may be particularly important for strand discrimination during MMR. Consistent with this notion, recent studies indicate that the ATPase function of MLH1/PMS2 is not required for nuclease activity *per se*, but appears important for maintenance of strand discrimination signals [51]. The precise role for ATPase function in the regulation of MLH1/PMS2 remains poorly understood, which as with all of the mutants described here, would be informed by additional structural information in the MMR complex combined with PCNA and RFC.

## METHODS

### Oligonucleotides, plasmids, and cell lines

The reporter plasmid 3%RMD-GFP was previously described [1]. The sgRNA/Cas9 plasmids used were px330 plasmids: Addgene 42230, deposited by Dr. Feng Zhang [55]. The sgRNA sequences for inducing DSBs in the reporter were previously described [1]. The plasmids pCAGGS-NZE-GFP (GFP expression vector), pgk-puro, and pCAGGS-BSKX empty vector (EV) were described previously (22941618). The WT expression vectors for MLH1 and PMS2, and PMS2-E702K, were previously described [11]. The mutant forms of MLH1 (KQQ, 566-3A, E34A, and RADA) and PMS2 (721-4A) were generated with gBLOCKs (IDT).

The WT and *Mlh1^-/-^* mESC lines with the RMD reporter previously described [1, 11]. The *Pms2^-/-^* mESC line was derived using two Cas9-mediated DSBs to introduce a deletion in *Pms2* using the following sgRNAs, cloned into px330: 5’ GACTTGGACTGCCGTCCTCC and 5’ CCGTGGCTCGCAGGACAAAT. WT cells were transfected with the corresponding plasmids and pgk-puro using Lipofectamine 2000 (Thermofisher), enriched for transfected cells using transient puromycin (Sigma Aldrich) treatment, then plated at low density to isolate and screen individual colonies for loss of PMS2.

### DSB Reporter Assays

For the RMD assays, mESCs were seeded at a cell density of 0.5 x 10^5^ cells per well of a 24-well plate, with 0.5 ml media. The next day, each well was transfected with 200 ng of each sgRNA/Cas9 plasmid using Lipofectamine 2000 (Thermofisher), with 0.5 ml of antibiotic free-media. For the RMD assays with expression vectors for various genes, transfections included 200 ng of those vectors, or the EV control (pCAGGS-BSKX). Each experiment had parallel control transfections using the GFP expression vector, along with the corresponding expression vectors, which was utilized to normalize all repair frequencies to transfection efficiency. For the RMD frequency assays, three days after transfection cells were analyzed by flow cytometry using ACEA Quanteon, as described [56].

### Mismatch Resolution Analysis

The mismatch resolution analysis was previously described [11]. Cells were seeded at a density of 0.1 x 10^6^ cells per well of a 12-well plate with 1 ml of media and transfected with the plasmids expressing the sgRNAs/Cas9 and relevant expression vectors. Three days after transfection cells were expanded prior to sorting for GFP+ cells, which were cultured for sorting a second time (BD Aria). Genomic DNA from each sample was then purified by phenol/chloroform extraction as described [56], then was used to amplify the repeat sequence using RMDjunct368UPillumina (5’ACACTCTTTCCCTACACGACGCTCTTCCGATCTCCGGGTCCTTCTTGTGTTTC) and RMDjunct368DNillumina (5’GACTGGAGTTCAGACGTGTGCTCTTCCGATCTAACAGCTCCTCGCCCTTG) primers, which include Illumina adapter sequences. The amplicons were subjected to deep sequencing and aligned the reads to the top strand sequence (Amplicon-EZ analyasis, AZENTA/GENEWIZ) (Fig 2A), reads that represented ≥0.1% of the total reads were individually aligned to the reference sequence, and each of the 8 mismatches were identified as being from either the top or the bottom strand (Fig 2A) and used to calculate the percentage of top strand base retention at each mismatch. Each cellular condition was examined with three independently transfected wells and GFP+ sorted samples and used to calculate the mean and standard deviation for each condition.

### Immunoblotting and quantitative reverse transcription PCR (qRT-PCR)

For immunoblotting, cells were transfected with the same total plasmid concentration as for frequency experiments but using EV instead of sgRNA/Cas9 plasmids, and were scaled 4-fold using a 6-well dish. Three days after transfection, cells were lysed using ELB buffer (250 mM NaCl, 5 mM EDTA, 50 mM Hepes, 0.1% (v/v) Ipegal, and Roche protease inhibitor) with sonication (Qsonica, Q800R). Blots were probed with antibodies for MLH1 (Abcam, ab92312), FLAG (Sigma, A8592), and ACTIN (Sigma, A2066), secondary antibody (ab205718). ECL reagent (Amersham Biosciences) was used to develop immunoblotting signals.

For quantitative RT-PCR (qRT-PCR) analysis to examine mRNA levels, total RNA was extracted using RNAeasy (Qiagen) and reverse transcribed using MMLV-RT (Promega). The RNA primers for PMS2 (5’ATAACGTGAGCTCCCCAGAA; 5’ GAGGACCAGGCAATCTTTGA) and ACTIN (5’GGCTGTATTCCCCTCCATCG; CCAGTTGGTAACAATGCCATGT) were used to amplify target mRNA using iTaq Universal SYBR Green (Biorad, 1725120) and quantified on (Biorad CRX Connect Real-Time PCR Detection System, 1855201). Cycle threshold (Ct) was used to determine relative levels of PMS2 mRNA subtracted by the Ct value of ACTIN for individual PCR samples (ΔCt value). The ΔCt value was then subtracted from the corresponding ΔCt from WT cells (ΔΔCt value), which was then used to calculate the 2^-ΔΔCt^ value.

